# Superbugs online: Co-production of an educational website to increase public understanding of the microbial world in, on and around us

**DOI:** 10.1101/2023.11.17.567568

**Authors:** Jon M. Tyrrell, Sarah Hatch, Melissa Flanagan, Kerry Owen, Yvonne Proctor, Catherine Stone, Geoff Fricker, Kirk Hullis, Matthias Eberl

## Abstract

1.

Digital tools and online presence have become a cornerstone to public engagement and involvement strategy and delivery. We here describe the co-production process behind launching a new multilingual resource for schools in the UK and beyond, jointly between university scientists, engagement professionals, primary and secondary teachers, and web designers. The ‘Superbugs’ website aims at raising awareness and increasing public understanding of the microbial world in, on and around us, and has attracted >19,000 online visitors, >33,500 page views and >775,000 Twitter impressions over the past 24 months. *Superbugs*.*online* is available in English, Welsh, Irish and Scottish Gaelic, thus making it accessible to everyone in the UK and Ireland, regardless of the language in which they receive and deliver their science education. The website is easy to navigate and features background information, quizzes, animations, videos, illustrated stories, interactive timelines, games and protocols for home experiments. All materials are presented in a non-prescriptive way, aimed at allowing flexibility for the materials to be adapted to the individual needs of teachers and pupils alike. Our work has led to demonstrable impact on the co-production team and on pupils and teachers as key stakeholders, based on a comprehensive evaluation of the co-production process itself, the impact of the end product, and the creation of lasting relationships with stakeholders and co-producers, for the mutual benefit of everyone involved.

**Data summary:** The authors confirm all supporting data have been provided within the article or through supplementary data files.

## 3. Introduction

Digital tools and online presence have become a cornerstone to public engagement and involvement strategy and delivery [1, 2]. The COVID-19 pandemic enforced an involuntary pivot to such virtual approaches – extended periods of lockdowns, closure of public spaces and schools, and social distancing meant that public engagement activities had to adapt, or stand still [3, 4]. ‘Superbugs’, a research-driven initiative at the universities of Cardiff and Swansea, aimed at improving public understanding of the microbial world in, on and around us [5], was no different. The original goal of ‘Superbugs’ was to investigate the mechanics of how to communicate and educate on complex topics such as microbiology and antimicrobial resistance (AMR), through novel engagement strategies in public spaces [5]. When it became impossible to conduct in-person outreach throughout most of 2020 and 2021 as a result of COVID-19 restrictions, the new aspiration was to widen the scope and catchment of communicative work around research and education, condense the enthusiasm from in-person events into an equally inspiring virtual format, and in partnership with teachers co-produce an interactive educational online resource.

The principle of co-production within this context may be most succinctly defined by the National Institute of Health Research (NIHR); *‘an approach in which researchers, practitioners and the public work together, sharing power and responsibility from the start to the end of the project’*, based on the assumption *‘that those affected by research are best placed to design and deliver it’* [6]. Central to this is the fact that the partners in the co-production process would also be the primary beneficiaries of its outcomes. In the case of Superbugs, we specifically wanted to target teachers and students of Key Stages 2 (KS2) and 3 (KS3) (school years 3-9 in England and Wales).

A co-production approach was particularly beneficial in the creation of the website as the project coincided with the launch of the new Curriculum for Wales [7]. As such, a key ambition of the Superbugs project was to develop educational resources that would meet the ‘four purposes’ that aim to support children and young people to be (i) *‘ambitious, capable learners, ready to learn throughout their lives’*; (ii) *‘enterprising, creative contributors, ready to play a full part in life and work’*; (iii) *‘ethical, informed citizens of Wales and the world’*; and (iv) *‘healthy, confident individuals, ready to lead fulfilling lives as valued members of society’*. Similarly, despite a primary focus on microbes, infection and antibiotics we wanted to ensure that our materials spanned across all six ‘areas of learning and experience’ defined as *Health and Well-being*; *Science and Technology*; *Mathematics and Numeracy*; *Expressive Arts*; *Humanities*; and *Languages, Literacy and Communication*. The expertise and experience of co-producing teachers ensured that our resources aligned with the content and the spirit of this new framework [7].

Additionally, in line with the Welsh Government’s aim to promote and facilitate the use of the Welsh language and to ensure that Welsh is treated no less favourably than English [8], we were keen to provide all Superbugs resources bilingually from the start. This was to take into account the fact that there is a free choice of Welsh and English-medium schools in Wales, and to make ‘Superbugs’ equally available to all children in Wales independent of the language in which they receive their education.

The Co-production Network of Wales outlines five key values that underpin successful co-production: (i) *‘value people and build on their strengths’*; (ii) *‘develop networks that operate as silos’*; (iii) *‘focus on what matters for the people involved’*; (iv) *‘build relationships of trust and shared power’*; and (v) *‘enable people to be change makers’* [9]. These values were kept central to our project at all times. Herein we provide an account of our co-production process, its implementation in launching a new microbiology resource for primary and secondary schools in Wales and beyond, the evaluation of both the co-production process itself for developing educational materials and the initial impact of the end product, and the creation of lasting relationships with stakeholders and co-producers, for the mutual benefit of everyone involved.

## 4. Methods

### Pre-production preparation

Three interactive sessions by the Co-production Network for Wales (https://www.copronet.wales) for the Superbugs core members (Tyrrell, Hatch, Eberl) and two professional web designers (Fricker, Hullis) emphasised the underlying concepts and philosophies of co-production, and how to design, implement and evaluate the process. Recruitment of co-production partners was primarily by utilising the existing communication network of all schools from across Wales, compiled by the Cardiff University School of Medicine Engagement Team. While being limited to virtual meetings due to COVID-19 restrictions in place at the time [10], and not being able to take advantage from in-person teambuilding as foundation for constructive co-production, we were able to involve teachers from the whole of Wales, thus covering a geographical area of 20,000 km^2^. These virtual meetings benefited from our extensive experience with the all-Wales ‘Life Science Challenge’ inter-school competition, which features online quiz rounds [4], and the development and delivery of online and hybrid teaching material both for school children and university students. We recruited 25 teachers to attend an initial introductory session, delivered over Zoom, where they were introduced to the background and philosophies behind the ‘Superbugs’ initiative, the idea of developing an educational tool for remote learning, and their potential role in this project. Teachers represented primary and secondary schools across Wales and covered both English and Welsh-medium education. 15 teachers expressed an interest to continue to be involved in the project and became our official co-production partners.

### Co-production process

Once partners had been recruited, co-production unfolded through three online workshops, delivered over Microsoft Teams and facilitated using interactive tools such as Mentimeter (www.mentimeter.com), Slido (www.slido.com) and Padlet (padlet.com). All co-production workshops were attended by university scientists and engagement professionals, primary and secondary school teachers, and web designers, to ensure that all scientific, educational and technical aspects of the project were represented. With the official launch of the Superbugs website (www.superbugs.online) in October 2021, the project entered the post-co-production phase.

### Evaluation of processes and outputs

As with our Superbugs pop-up shop project [5], a multifaceted evaluator approach was taken. This was in part directed by the use of the ‘Measuring What Matters’ tool, provided by Co-production Network for Wales (info.copronet.wales/audit-online), a practical task which allowed us to identify the important questions to ask in order to rigorously achieve our evaluation aims. As a result, it was agreed that our evaluation would consist of four distinct elements.

The <u>horizontal evaluation</u> comprised of a self-assessment by project participants combined with peer review, primarily via a self-audit at the beginning, during, and at the end of the project.

For the p<u>articipatory approach evaluation</u>, we collected direct feedback from primary stakeholder participants, through bespoke questionnaires for teachers and school children during the ‘Piloting Draft’ element of the project.

The <u>impact evaluation</u> consisted of two parts – an evaluation of the impact upon our co-production partners with regard to their own personal knowledge of microbiology and AMR, and their self-reported confidence and competence in teaching on these topics.

Finally, for the <u>product evaluation</u> we assessed the success of the Superbugs website itself, gauged through monitoring website traffic, feedback collected through questionnaires and interactive elements within the website, and anecdotal feedback.

## 5. Results and Evaluation

### Outcomes of co-production workshops

The overall study design, including a series of co-production workshops, is visualised in **Figure 1**. When conceiving the project, an initial logic model was developed during the pre-production phase to define our aspirations and the processes by which we would achieve our aims and objectives. Upon recruitment of our co-production partners, the logic model evolved further to more accurately reflect the changing focus and priorities of the wider team. This revision formed a repeated exercise through the course of three co-production workshops, the final output of which can be seen in **Figure 2**.

**Figure 1.**
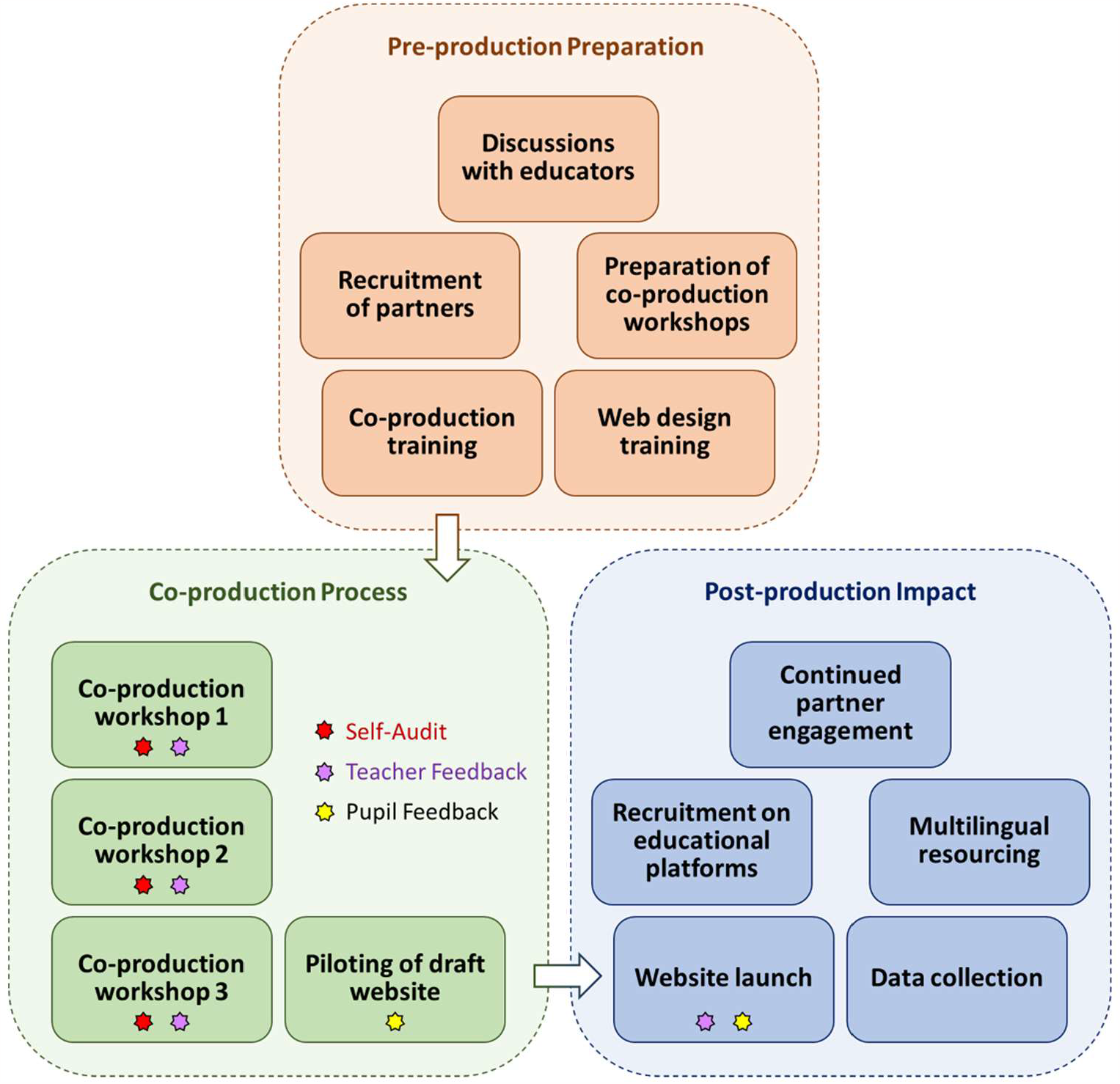
Schematic overview of the co-production project structure. Time line: Pre-production preparation, November 2020 – February 2021; Co-production process, March 2021 – September 2021; Post-production impact, October 2021 – September 2023.

**Figure 2.**
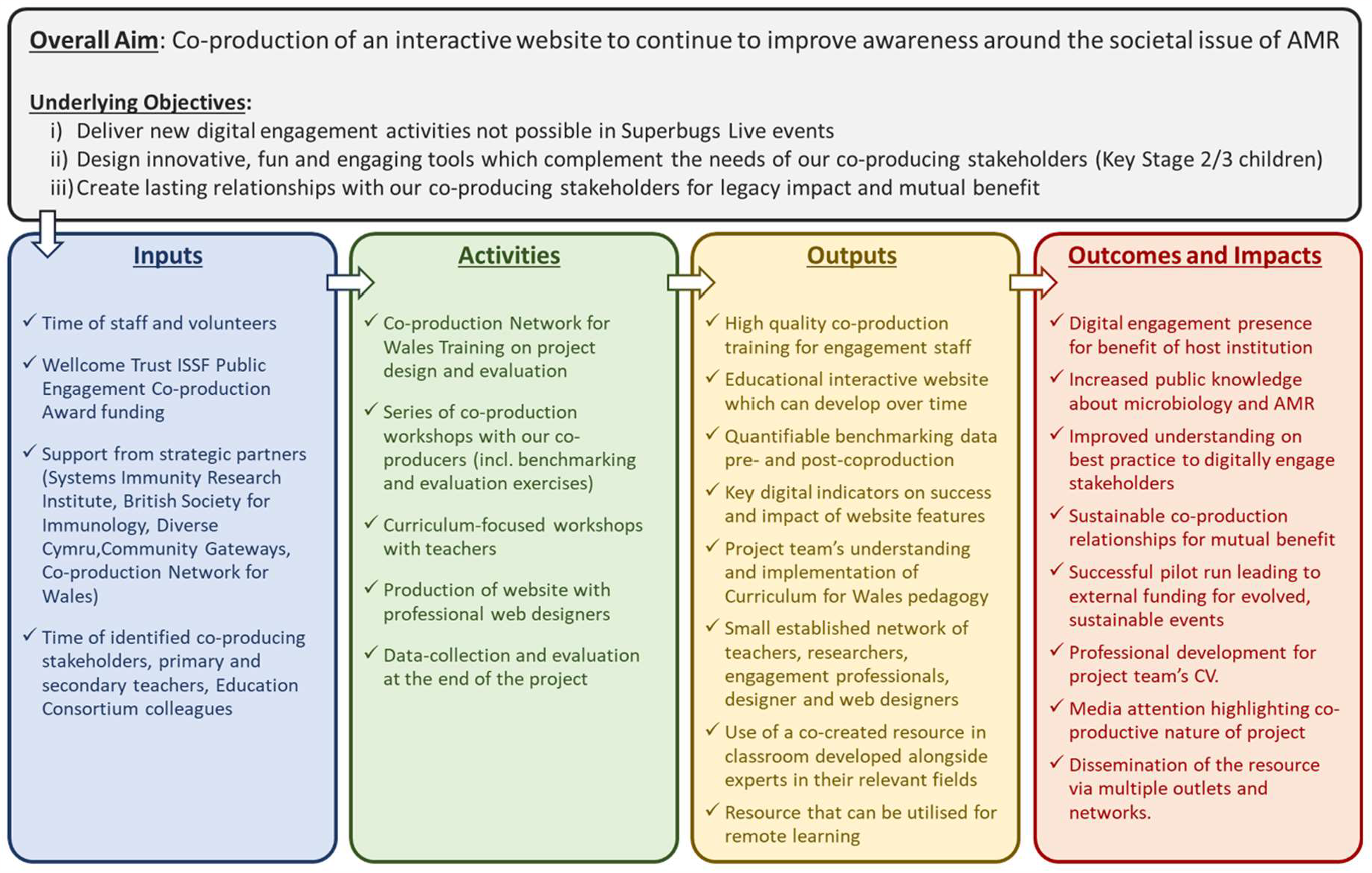
Co-produced logic model illustrating the relationship between project resources, activities and intended impact. This is based on an initial draft that was amended and expanded during the co-production workshops.

<u>Workshop 1</u> was an in-depth introduction to the project and what we were hoping to achieve. It provided a useful understanding of teachers’ individual experience in designing, using and evaluating digital teaching during the COVID-19 pandemic when schools were forced to resort to remote and hybrid learning over extended periods – a setting which was entirely novel for both teachers and pupils at the time (**Table 1**). In particular, insight was gained into a range of educational online platforms already employed, and how best to communicate for the remainder of the project (**Figure 3A**). These formed the basis of our delivery strategy for the pilot development of *Superbugs*.*online*. Teachers also undertook a benchmarking exercise in form of a simple quiz, delivered via Slido, to gauge their level of knowledge of basic microbiology.

**Table 1:**
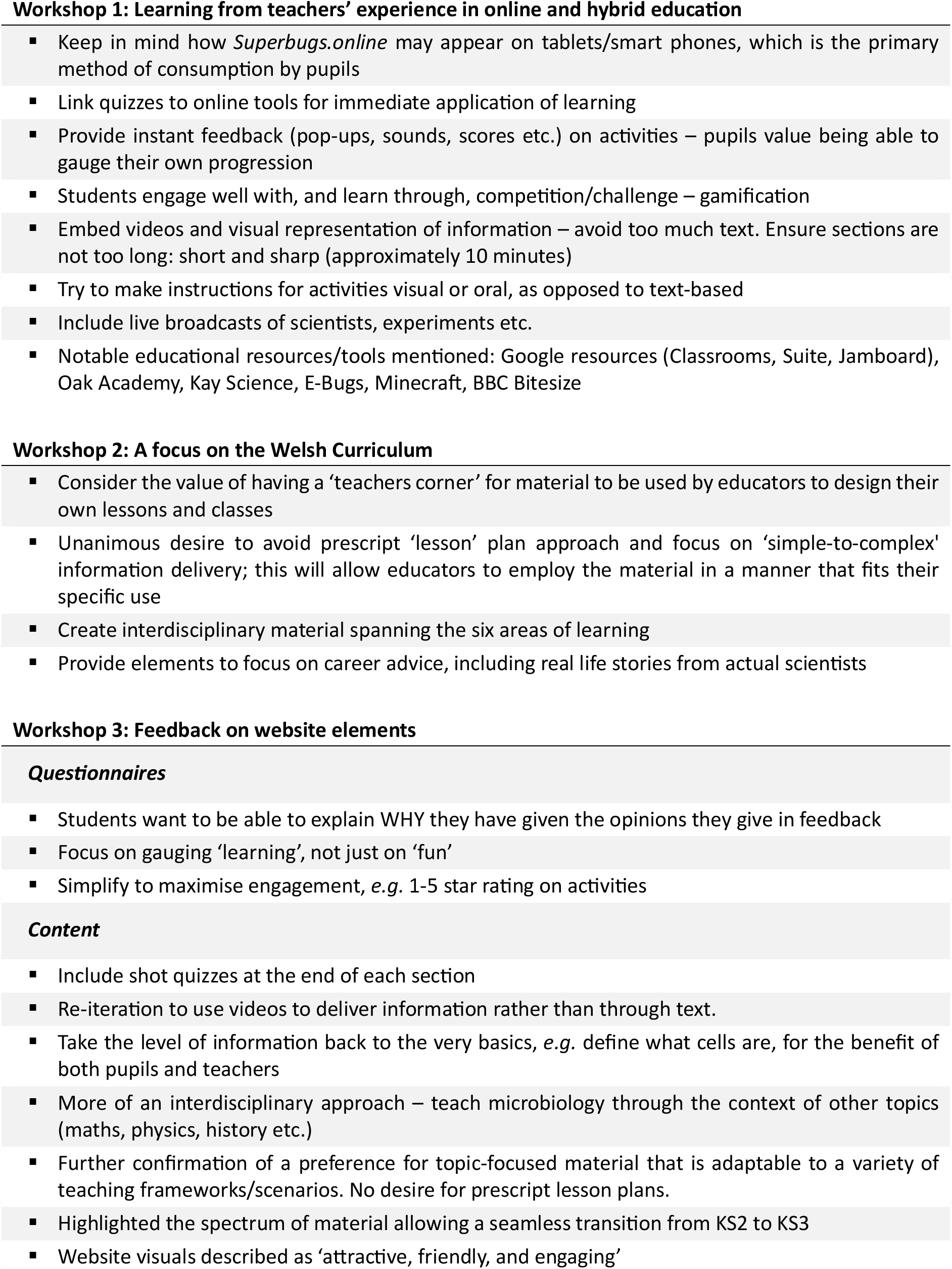
Significant outputs and discussion points from the co-production workshops.

**Figure 3.**
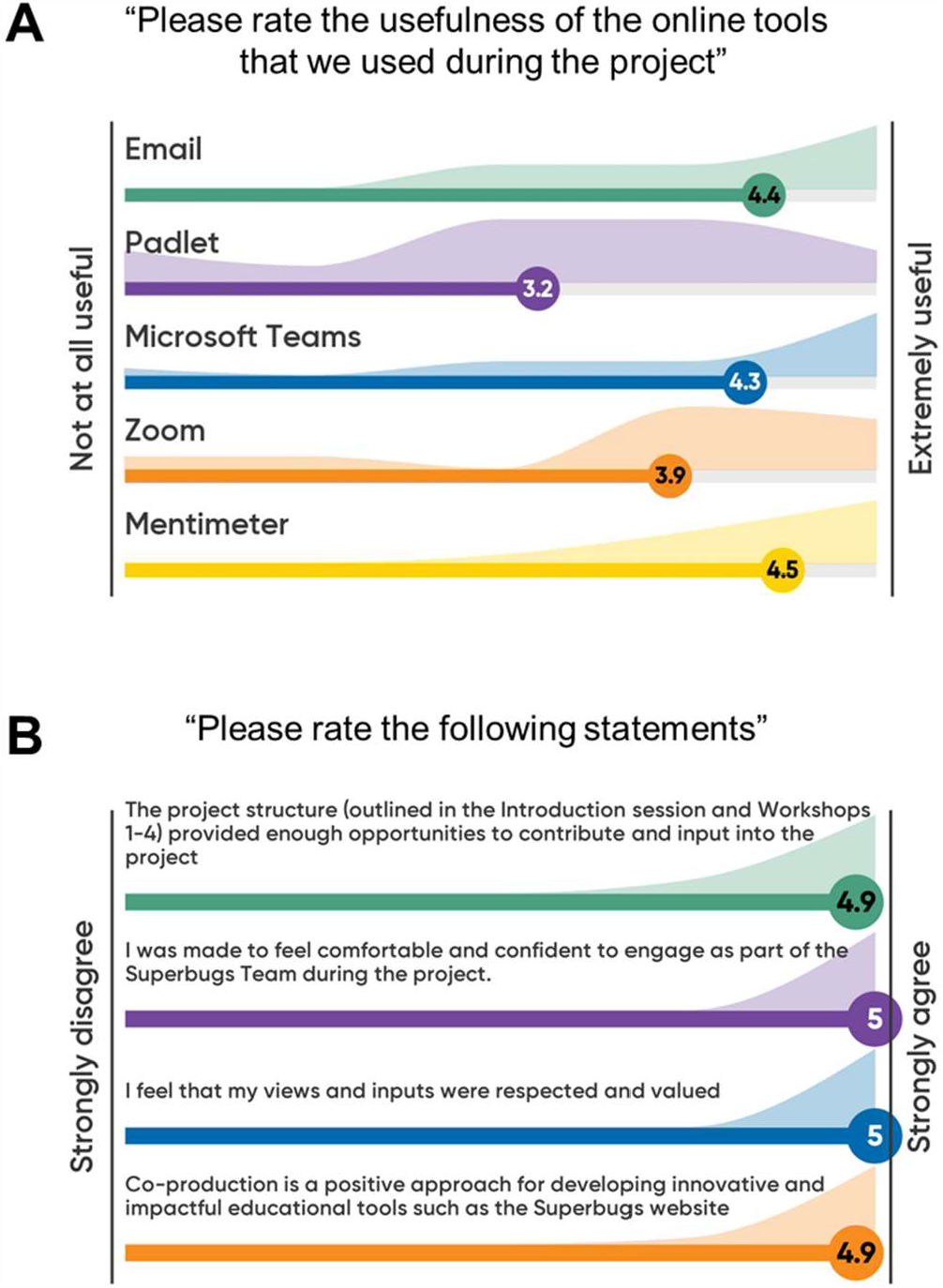
Evaluation of communication methods via electronic means and self-reflection of teachers on the co-production process. Feedback from 14 primary and secondary school teachers involved in co-producing the Superbugs website was collected during the co-production workshops using Mentimeter. Possible answers and scored ranged from 1: “*Not at all useful*” to 5: “*Extremely useful*” ***(A)***, and from 1: “*Strongly disagree*” to 5: “*Strongly agree*” ***(B)***.

<u>Workshop 2</u> was centred around developing *Superbugs*.*online* in close alignment with the needs of our educational partners, and was split into two parts. Firstly, we delivered an interactive lecture, introducing teachers to key concepts of microbiology, infection and AMR. This ensured all participants were on a consistent level of knowledge for the remainder of the project and able to engage constructively with the scientific aspects of the discussions. The second half of the workshop included discussions around the new Curriculum for Wales [7], to determine how our ambitions might fit into this new scheme, and whether and how its confinements might affect the design and content of *Superbugs*.*online*. A key point raised unanimously by all teachers was the preference for a non-prescript, flexible resource which would allow educators to apply and adapt *Superbugs*.*online* for their own purposes. To quote our partners directly; “*Prescription takes away from being able to adapt material to variety of ELOs [expected learning outcomes] across the curriculum*”, and “*having golden nuggets [of microbiology] that people can incorporate into their own teaching*… *would be more useful*”.

In <u>Workshop 3</u> we began to develop the initial pages of the website to be reviewed and discussed, and co-produced feedback questionnaires that would sit across the Superbugs platform, to allow for continuous monitoring of usage and feedback. Pupil feedback questions were co-produced with the help of a year 5 class at Llanedeyrn Primary School, Cardiff, and year 11 pupils at Ysgol Clywedog, Wrexham, with support by their teachers. Additionally, we collected valuable feedback from teachers on the pilot version of *Superbugs*.*online* and on how best to promote this website (**Table 2**).

**Table 2.** Suggestions by teachers on how best to promote our educational resources throughout schools in Wales. Feedback from 14 primary and secondary school teachers during the co-production phase of the Superbugs website.

| Suggestion | Action taken |
| --- | --- |
| *"Pencils and mugs with web address into staff rooms. Teachers love pencils and mugs."*<br>*"Merchandise"*<br>*"Superbugs-branded chocolate biscuits"* | Purchase of Superbugs-branded merchandise such as water bottles, bookmarks, pencils, torches and rulers that are given out at events |
| *"Are there events you can attend where all schools meet?"*<br>*"At events where all schools meet."*<br>*Are there teaching meetings/conferences that we can showcase at?* | Distribution of brochures at Cardiff University's 'Science in Health Live!' event 2023 and at the Teacher and Career Adviser's Conference 2023 |
| *"Social media – Twitter."*<br>*"Twitter, Facebook groups"*<br>*"Facebook. Email. Hwb."* | Establishment of a very active Twitter profile; also creation of accounts on YouTube and TikTok |
| *"Get in touch with Hwb.gov.wales and ask them to include it."*<br>*"Facebook. Email. Hwb."* | *Superbugs.online* now hosted on the Hwb resources page of the Welsh Government and on the 'Gaelic Education in Scotland' page hosted by Stòrlann |
| *"Email out to science co-ordinators"* | Targeted campaign to reach science teachers and/or science departments via direct messaging on Twitter |
| *"Use teachers communications network?"*<br>*"Through the network we have developed."* | Successful contacts with science teachers across Wales and beyond via Email and Twitter; establishment of long-term relationships with some of the co-production partners |

### Design of the Superbugs website

As a direct result of the co-production workshops we created a website aiming to be informative, self-explanatory and non-prescriptive, and at the same time easy to navigate, interactive and fun to visit (see Supplementary Information). Careful consideration of software solutions and design options ensured that the final product was straight-forward to maintain and develop further by the Superbugs team, without need for extended support by professional web designers and programmers after the initial set-up and training. *Superbugs*.*online* was launched bilingually in English and Welsh in October 2021 (**Supplemental Figure S1, Supplemental Table S1**). The ease with which content could be hosted in different languages quickly made us realise the full potential of the project, and with the help of Stòrlann Nàiseanta na Gàidhlig we were able to launch a Scottish Gaelic version in 2022. Grant funding from An Chomhairle um Oideachas Gaeltachta agus Gaelscolaíochta (COGG) enabled us to have an Irish version in place ready for the new school year 2023/2024. Thus, *Superbugs*.*online* now covers all four languages officially used for education at public and private schools across the British and Irish Isles. With a paucity of high-quality modern teaching resources in Welsh, Gaelic and Irish (compared to English), especially in STEM subjects, this puts the Superbugs project in a unique position to fill an urgent educational need and meet the demands and ambitions from teachers and pupils [11, 12].

### Impact on co-production partners

At the end of the co-production process, the partners were asked to undergo a self-evaluation and provide feedback on individual aspects of the project. All teachers (100%) among the co-production team agreed or strongly agreed with the notion that the project structure had provided enough opportunities for them to contribute, that they had been made feel comfortable and confident to engage, that they had felt their views and inputs were respected and valued, and that co-production was a positive approach for developing innovative and impactful educational tools (**Figure 3B**). Importantly, all teachers (100%) also considered that the level of their involvement had been just right for them, with nobody feeling that the level had been too little or too much (data not shown). This positive evaluation was complemented by constructive and supportive verbal feedback (**Table 3**), demonstrating that the co-production element had been designed optimally to benefit from involving key stakeholders in the conception, design and implementation of the project.

**Table 3.**
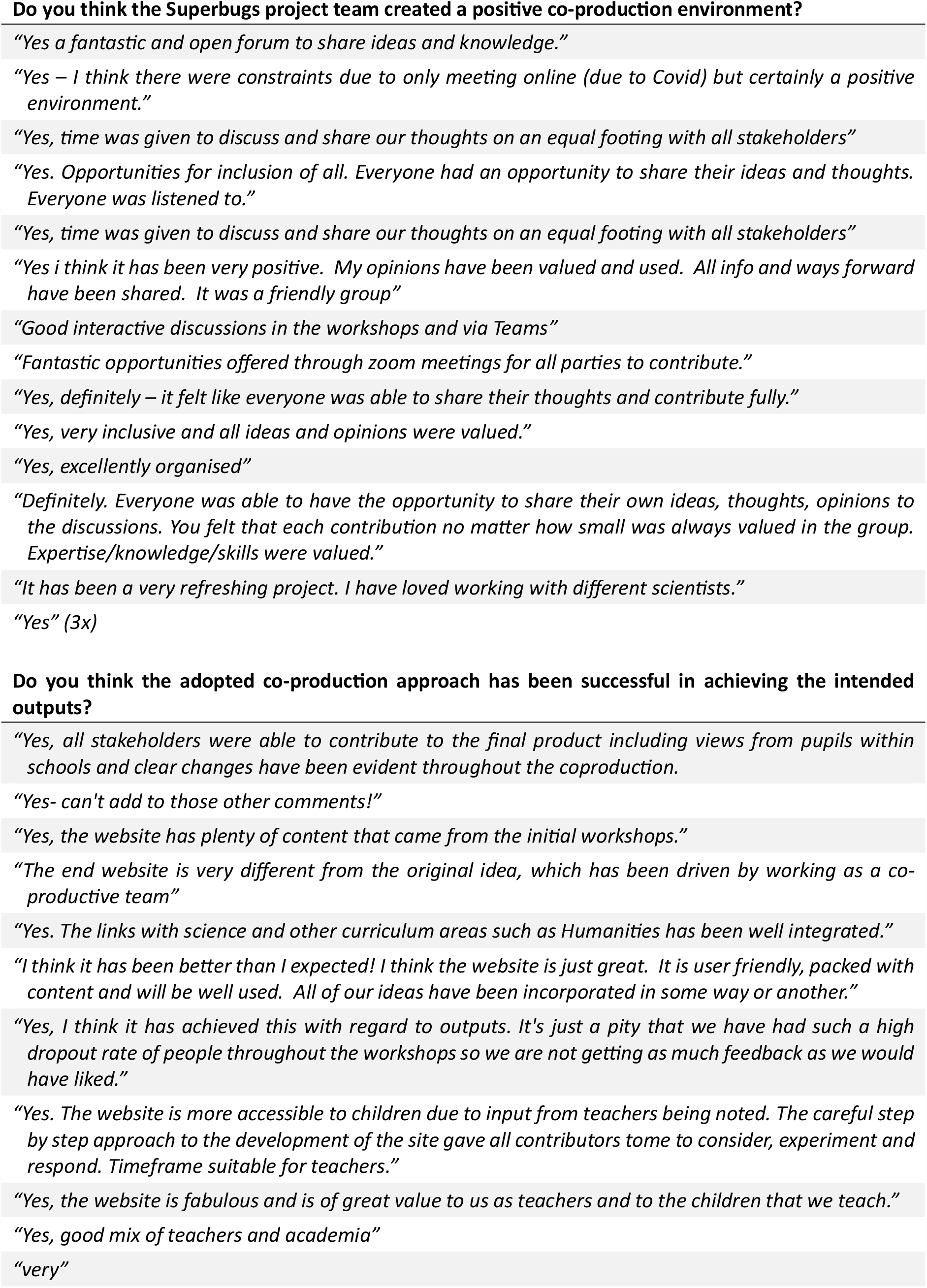
Self-reflection of teachers on the co-production process. Feedback from 14 primary and secondary school teachers involved in co-producing the Superbugs website.

The Superbugs core team together with the web designers completed the Co-production Network for Wales’s self-audit tool before, during and at the end of the project. At the beginning of the projects, all five categories (Assets, Networks, Outcomes, Relationships, Catalysts) only ranked either as “Made a start” or “Making progress” (**Figure 4**). This initial assessment had improved considerably by the end of the project, with all five categories either ranking as “Doing well” or “Doing as well as you can”, as testimony of the achievements and learnings during the co-production phase.

**Figure 4.**
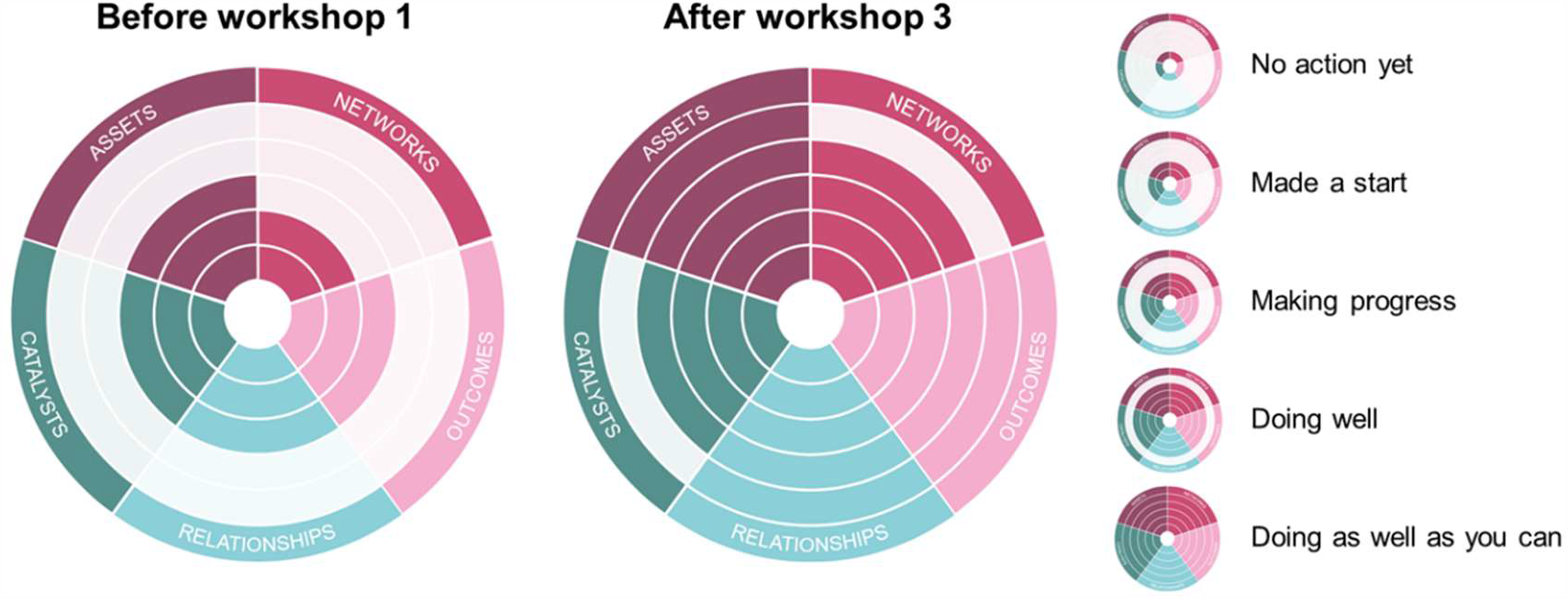
Outcomes of the self-audit at the beginning and at the end of the co-production process. Definitions: ASSETS, *“You value all participants and you build on their strengths and resources*.*”* NETWORKS, *“You develop networks of mutual support*.*”* OUTCOMES, “*You do what matters for all people involved with a focus on outcomes*.*”* RELATIONSHIPS, *“You build relationships of trust, reciprocity and equality by sharing power and responsibility*.*”* CATALYSTS, *“People are change makers, as an organisation your role is to enable this*.*”* Steps from the centre of the diagram to the periphery: No action yet; Made a start; Making progress; Doing well; Doing as well as you can. This tool was developed by the Co-production Network for Wales (https://info.copronet.wales/the-self-evaluation-audit-tool).

### Impact on participating teachers

It was important to the outcomes of the project that we imparted a positive impact on all co-production partners. As such, during Workshop 1, a simple benchmarking exercise was carried out to ascertain the baseline knowledge of our partners on topics around microbiology, AMR and antibiotic stewardship. All teachers (100%) showed a basic working understanding of antibiotics, their selective toxicity against micro-organisms, and some basic good stewardship practice. This was encouraging as being informed to this level would enhance the intellectual level of input that could be gained throughout the co=production phase. However, many teachers suggested to have reservations in their ability to communicate/educate these topics further, with only 46% feeling confident enough to deliver teaching on AMR and microbiology. Additionally, 31% of teachers indicated at the beginning of the project that they did not feel informed enough to teach on these topics, compared to 69% who answered “a little bit” or “very much”. Importantly, this original hesitation quickly disappeared so that by the end of the project all teachers (100%) felt informed and confident in their abilities to design and deliver classes framed around the content of *Superbugs*.*online*.

### Impact on pupils

During the development of *Superbugs*.*online*, early versions of the website were shared with pupils, and feedback from a total of 247 pupils was received. Despite only accessing a prototype of our resources, it was reassuring that 78.1% of pupils rated the preliminary website ‘excellent’ or ‘good’, and that 38.0% would have been happy to share it with friends (compared to 11.4% who would not be happy to do so) (**Supplemental Figure S2**). Verbal feedback emphasised the breadth of content that the students found interesting (**Supplemental Table S3**).

From the time of the official launch, student feedback was collected via feedback forms embedded in the Superbugs website. To avoid biases due to responses from individual visitors with a markedly positive or negative view of our resources, we here focused on feedback given by whole groups of students accessing the website jointly. In one particularly informative exercise at the time of the launch, all students of the same year 6 class attending an English-medium primary school in Cardiff were encouraged by their teacher to complete the online feedback form. Out of 22 submitted questionnaires, 59.1% of pupils rated *Superbugs*.*online* ‘excellent’ or ‘good’, 72.7% found it excellent or good navigate, and 68.2% found the graphics and visuals excellent or good (**Figure 5A**).

**Figure 5.**
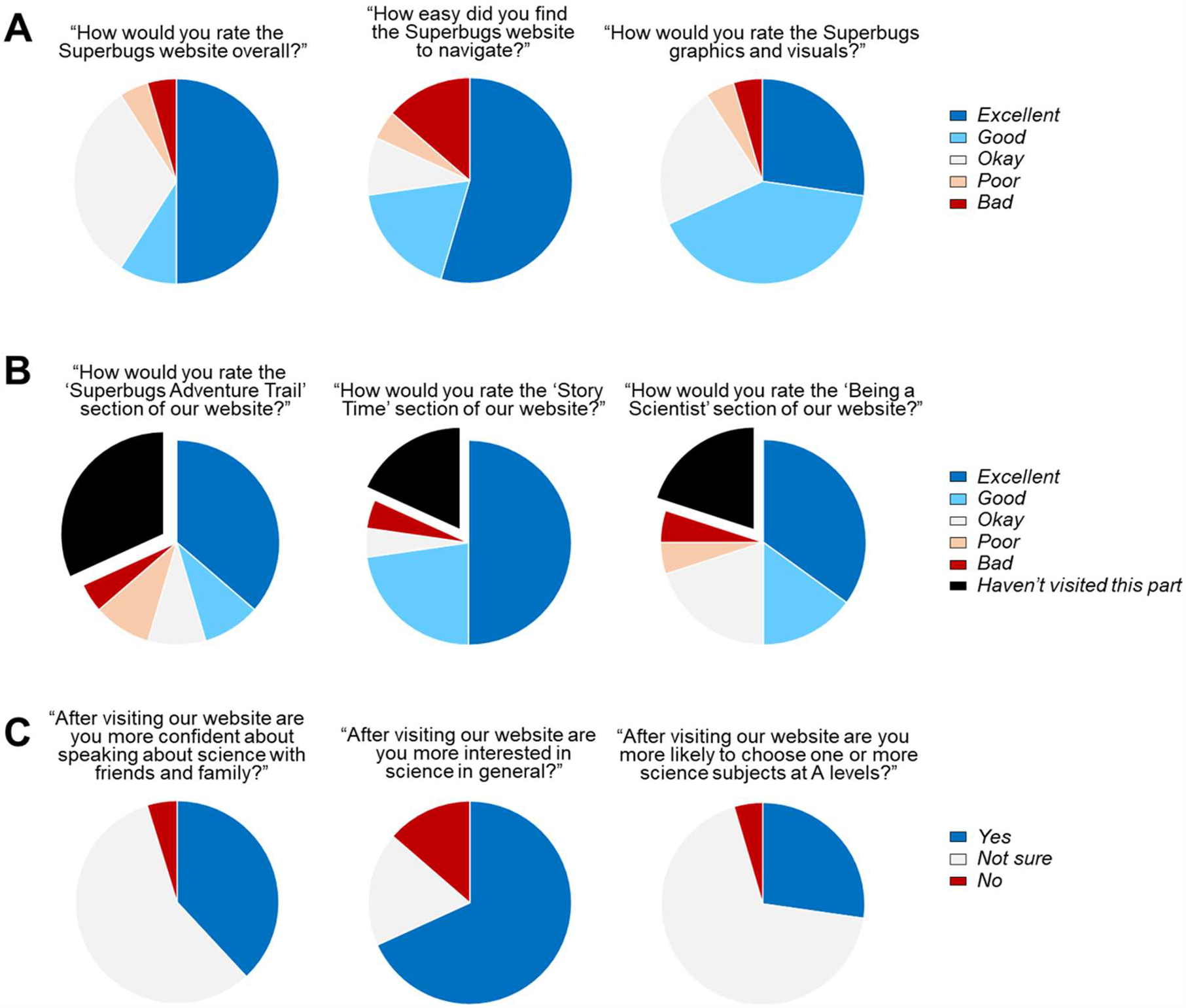
Feedback from pupils attending an English-medium primary school in Cardiff. ***(A)*** Overall website rating. ***(B)*** Ratings of major website sections. ***(C)*** Personal experience. Data were compiled from the answers given by 22 pupils of the same year 6 class completing the online feedback form in October 2021, shortly after the launch of *Superbugs*.*online*.

Of the pupils who visited the corresponding sections, 66.7% rated the ‘Adventure Trail’ excellent or good, 88.9% the ‘Story Time’ section, and 62.5% the ‘Being a Scientist’ section. Encouragingly, only very few pupils considered *Superbugs*.*online* overall or aspects of it ‘poor’ or ‘bad’ (**Figure 5B**), and at least one of these negative responses may have been due to a mistake as it was accompanied by positive comments in the free text parts of the questionnaire (not shown). After visiting *Superbugs*.*online*, 38.1% of pupils felt more confident about speaking about science with friends and family, 68.2% were more interested in science in general, and 27.3% stated they were more likely to choose one or more science subjects at A levels (the school leaving certificate in England and Wales); again, only very few pupils gave negative responses (**Figure 5C**). These overall positive impressions from the questionnaires were reinforced by anecdotal comments provided by pupils about the most interesting thing they had learned and their favourite part of *Superbugs*.*online* (**Supplemental Table S3, Supplemental Table S4**).

## 6. Lessons Learned

### Co-production

The co-production process led to the development of lasting relationships. Several partners remained with the project in a productive and cross-disciplinary team, even after the end of the official project, and are still being consulted on many aspects of *Superbugs*.*online*, and the wider Superbugs initiative. We very much see these partners now as a part of the core Superbugs team, and as such no longer officially evaluate their input. Whilst perhaps idealistic, this has certainly been an organic route of travel, and testifies to the long-term success of our co-production.

### Multilingual challenges

With its provision of multilingual materials, *Superbugs*.*online* meets a key educational need in the UK’s devolved nations (Wales, Scotland and Northern Ireland) and in the Republic of Ireland. However, accurate provision of educational content in four different languages has turned out to be more demanding than anticipated as it needs to be cross-checked by translators and education experts to make sure the texts are both scientifically accurate and at an appropriate language level for pupils [11-14]. It is important to note that some Superbugs content is at present provided in English only – *e*.*g*. certain videos, animations and online games where simultaneous translation is technically not possible. Ongoing work aims at replacing those English materials with multilingual alternatives and/or appropriate subtitles wherever possible. The inclusion of a glossary that lists scientific terms and explains them in simpler words for each language is planned but has not been started.

### Promotion

Creating an educational resource is only the beginning of raising awareness and increasing understanding as it needs to be promoted heavily to make it visible and stand out against potential competitors [15]. As such, social media campaigns, direct contact with teachers, a printed brochure and word of mouth recommendations were all important in promoting *Superbugs*.*online* and in ensuring its relevance for science education (**Supplemental Figures S3 and S4**). The co-production process helped considerably in this regard, allowing us to gauge the wishes and interests of teachers and pupils alike, whilst simultaneously developing materials that directly catered for our stakeholders’ needs. As of October 2023, *Superbugs*.*online* was already listed as a recommended educational website by Hwb (Digital Learning for Wales, the Welsh Government’s provision of educational tools), Gaelic Education (maintained by Stòrlann Nàiseanta na Gàidhlig), and the Federation of European Microbiological Societies (FEMS), with evidence of direct traffic referrals from each site.

### Feedback

Collecting meaningful feedback from anonymous visitors remains a challenge. Although *Superbugs*.*online* features three bespoke (and co-produced) questionnaires for pupils, teachers and other members of the public alike, uptake of this option has been very poor so far outside a supervised setting in the classroom. This is likely to be due to general survey fatigue [16, 17] and needs to be addressed going forward. The long-term aim is to embed more seamless evaluation strategies as part of the engaging activities. In the meantime, we will continue and intensify our collaborations with teachers and schools to learn how to improve our resources further, and keep monitoring web traffic to understand which parts of *Superbugs*.*online* are particularly successful. The web and Twitter statistics so far, together with direct feedback from teachers and pupils, demonstrate the effectiveness of our engagement with key stakeholders and the interest in our resources, thus directly addressing the original objectives set out in the co-production workshops.

### Long-term investment

Developing and debugging an educational resource such as *Superbugs*.*online* and keeping it up to date comes with considerable expenses, even if the content itself is largely contributed free of charge. Costs for long-term website maintenance currently amount to approx. £750 per year for domain registration and Squarespace and Weglot subscriptions, in addition to translation and proof-reading services. So far, our team has been able to cover those expenses from grants related to the development and translation of *Superbugs*.*online* but going forward this will require ongoing support to ensure a stable provision of our educational website in the long run.

### Added value for in person activities

The possibility to provide online materials to complement our in-person events is an attractive option for extended engagement with the public and was successfully explored at small Superbugs workshops held during the 2023 summer holidays at Swansea Central Library (Swansea, UK) and the Broadlands Fun Day (Bridgend, UK). In addition to our ever popular activities such as a microscope station and a ‘Grow your own Microbe’ body swab station, these events also featured guided tours through *Superbugs*.*online* and the possibility for visitors to explore the online content at their own pace. The ‘Behind the Scenes’ blog section now regularly contains summaries and photos from live events, thereby increasing the audience reach and promoting in-person activities and website alike.

## 7. The Future of Superbugs

Our approach of combining public engagement and co-production with analyses into the effectiveness of the delivery and the impact lends itself to further academic training and investigation. In this regard, we have started to offer public engagement-based projects for students of the MSc Biomedical Science (Clinical Microbiology) module at Swansea University (UK). The first four projects successfully completed addressed the development and assessment of novel in person activities or online content, in co-production with teachers and pupils, and comprised a pre-production phase, a pilot phase in a public setting, and the final delivery at a local school (Cefn Glas Infant School, Bridgend, UK). We believe that this is a powerful way to train the next generation of scientists and equip them with a rich set of transferrable skills in science communication, education, teamwork, creativity and evaluation.

AMR is of global concern, and improving modern science education and keeping it current and relevant is an ambition everywhere. The English content of *Superbugs*.*online* is naturally accessible to a worldwide audience, as reflected in the web traffic to our website from Europe, the Americas, Africa, Asia and Australia (**Supplemental Figure S5**). Building on this positive momentum, we are increasingly working with international partners raising awareness in their communities to explain the underlying scientific and health principles of hygiene, infections, vaccines and AMR, for instance with volunteers in Tanzania and Liberia. We also participated in the virtual ‘Night of Science’ 2022 (bilingually in English and Ukrainian), organised by colleagues at Zaporizhzhia Polytechnic National University, Cardiff’s official partner university in Ukraine. These collaborations involve promoting and supporting local activities, participating in events and activities, and developing joint outreach programmes. Our Superbugs initiative will continue to drive improved microbial literacy worldwide through a growing portfolio of research-driven, innovative public engagement projects.

## 10. Author statements

### 10.1. Author contributions

J.T.: conceptualisation, funding acquisition, methodology, project administration, resources, data collection, formal analysis, writing – original draft, writing – review and editing. ORCiD 0000-0001-8565-2590 @JTyrrell_Micro. S.H.: conceptualisation, funding acquisition, methodology, project administration, resources, data collection, formal analysis, writing – original draft, writing – review and editing. ORCiD 0009-0008-9522-7553 @SGlenysHatch. M.F.: methodology, resources, data collection, writing – review and editing. K.O.: methodology, resources, data collection, writing – review and editing. Y.P.: methodology, resources, data collection, writing – review and editing. C.S.: methodology, resources, data collection, writing – review and editing. G.F.: methodology, writing – review and editing. K.H.: methodology, writing – review and editing. M.E.: conceptualisation, funding acquisition, methodology, project administration, resources, data collection, formal analysis, writing – original draft, writing – review and editing. ORCiD 0000-0002-9390-5348 @EberlLab.

### 10.2. Conflicts of interest

Geoff Fricker and Kirk Hullis are digital consultants and web designers, and received payments for their work on this project. All other authors declare that there are no conflicts of interest.

### 10.3. Funding information

The development and dissemination of S*uperbugs*.*online* received financial support from the Wellcome Trust ISSF3 scheme, Cardiff University’s Systems Immunity Research Institute, Cardiff University School of Medicine, the British Society for Immunology, and An Chomhairle um Oideachas Gaeltachta agus Gaelscolaíochta (COGG). This project was also boosted by winning Cardiff University School of Medicine’s Staff Appreciation And Recognition (STAR) Award 2021 for ‘Outstanding Contribution to Engagement Activities’ (J.M.T.), the Best Poster in ‘Education & Public Engagement’ at the Annual Congress of the British Society for Immunology in Liverpool 2021 (M.E.), and the Microbiology Society’s Microbiology Outreach Prize 2022 (J.M.T.).

### 10.4. Ethical approval

This study did not classify as research involving human subjects, human material or human data, and as such did not require approval by an appropriate ethics committee. The individuals involved in this project were public involvement representatives and not participants in a research study. All individuals provided verbal consent to take part in the discussion groups.

## 10.5 Acknowledgements

This project would not have been possible without the insight, help and support by numerous teachers and their pupils giving advice and feedback before and after the launch of S*uperbugs*.*online*, with special thanks to our co-production partners Clare Bowen (St Albans Roman Catholic High School, Pontypool), Meirion Callaghan (Deri View Primary School, Abergavenny), Alison Hughes (Trowbridge Primary School, Rumney), Connor Llewellyn (St Richard Gwyn Catholic High School, Barry), Gemma Mabbett (Pennard Primary School, Swansea), James Morgan (Birchgrove Primary, Swansea), Matthew Oliver (Bryn Bach Primary School, Tredegar), Simon Phillips (Idris Davies School 3-18, Abertysswg), Jenny Richards (Glais Primary School, Swansea), Rachel Stephens (West Monmouth School, Torfaen), and Samantha Taylor (Pontypridd High School, Pontypridd). We would also like to thank Christie Conlon, Ruth Lewis, Erin Nash and Mike Roberts for help with communication and design, Karen Edwards for facilitating contacts with schools across Wales, Gail Thomas for advice on data safety, Laine Skinner for creating a multilingual Wordle, Jordan Mathias for contributions in the early stages of the project, the Welsh Government’s Hwb team and the Central South Consortium for their endorsement, and the people behind the helpdesks of Squarespace, Weglot and Knight Lab for their quick and constructive advice every time we needed it. Special thanks go to the Welsh translation unit at Cardiff University and Tudur Jones for the translation into Welsh, Morag Maclean and everyone at Stòrlann Nàiseanta na Gàidhlig for the translation into Gaelic, and Jack O’Driscoll, Máire McCafferty, Jenny Mannion and Cathal Ormond for the translation into Irish. Our biggest shoutout goes to the thousands of visitors of our website, and to everyone interested in learning more about the microbial world in, on and around us. We developed these pages for you and hope you find them useful and inspiring.

## Abbreviations

AMR: antimicrobial resistance
STEM: Science, Technology, Engineering and Mathematics

## 12. Supplementary information

See online supplement.

## Notes

https://www.superbugs.online

