## Supplementary Information for "Superbugs online: Co-production of an educational website to increase public understanding of the microbial world in, on and around us"

*[1]Institute of Life Science, School of Medicine, Swansea University, Swansea, UK; [2]Public Involvement and Engagement Team, School of Medicine, Cardiff University, UK; [3]Ysgol Clywedog, Wrexham, UK; [4]Windsor Clive Primary School, Cardiff, UK; [5]Tredegarville Church in Wales Primary School, Cardiff, UK; [6]Llanedeyrn Primary School, Cardiff, UK; [7]Maxim Consulting Services, Bristol, UK; [8]Division of Infection & Immunity, School of Medicine, Cardiff University, Cardiff, UK; [9]Systems Immunity Research Institute, Cardiff University, Cardiff, UK*

### Supplementary information

#### *Web design and software solutions*

*Superbugs.online* was created on the Squarespace (www.squarespace.com) website building platform, using Weglot (www.weglot.com) to host multilingual content simultaneously in English, Welsh, Scottish Gaelic and Irish. The original setup of the domain and the website mechanics was facilitated by Maxim Consulting (Bristol, UK), with all further development and maintenance as well as creation of content done by the Superbugs core team. Interactive timelines were created using the TimelineJS tool (timeline.knightlab.com), interactive quizzes using FlexiQuiz (www.flexiquiz.com). Feedback data, uploaded files and other information received from visitors were stored on a secure OneDrive folder hosted at Cardiff University, using automated workflows for data transfer via Zapier (www.zapier.com). Site traffic was monitored using the inbuilt Squarespace functions and Google Search Console; data privacy was ensured by disabling Squarespace analytics tracking for visitors who chose not to accept the corresponding analytics cookies, and by providing a detailed privacy notice on the website as approved by the compliance and risk advisors at Cardiff University responsible for data protection. To create and promote website content via social media, *Superbugs.online* was linked to profiles on Twitter (@CUsuperbugs), TikTok (@prof_superbugs) and YouTube (@superbugs4995).

#### *Major features of the Superbugs website*

The step-by-step 'Adventure Trail' of *Superbugs.online* features a wide range of engaging activities and information, consisting of (currently) 19 different topics. It starts with a basic introduction to life, evolution and bacteria, proceeds to harmless commensals, microbes in the environment and hygiene, and leads to the concepts of infection, antibiotics and vaccines. These topics are covered via different means such as introductory texts, images, animations, videos, timelines, games and quizzes. Importantly, visitors are free to explore the topics of the adventure trail either in the suggested order, or access selected themes directly, depending on individual interests – or perhaps as guided by a teacher.

The 'Adventure Trail' links to various complementary sections including a 'Downloads' section for additional information, colouring sheets and protocols for safe experiments to be conducted at home and in the classroom. Our 'Story Time' reading corner features illustrated stories about major breakthroughs in the scientific and clinical understanding of infections and how to fight them. In the 'Being a Scientist' section visitors can explore the types of jobs people working *in science* actually do, across a wide spectrum of scientific careers – be it as lab researcher, data scientist, clinician, teacher, science journalist, research funder or engagement professional – and what inspired these people to choose a career in science. The 'Who We Are' section presents the people behind Superbugs and features a 'Behind the Scenes' blog with personal insights from the team, news and updates, and contact details on how to get in touch and how to provide feedback. The 'Superbugs – Live!' section summarises our experience with in-person activities such as live pop-up shops in the city centres of Cardiff in 2019 and Exeter in 2023, link back to the 'Adventure Trail'.

Other noteworthy features of *Superbugs.online* include a 'Teacher's Corner' with practical information aimed at science teachers interested in using our resources in their classes; a 'Hall of Fame' to display artwork and documents produced by our visitors; a 'Games' corner including a multilingual science-focused Wordle puzzle available in English, Welsh, Gaelic and Irish; a virtual 'Bookshelf' with recommended popular science books expanding on topics explored on the Superbugs website; a 'Science Fun in Wales and Beyond' section showcasing events and activities open to anyone interested in biomedical science and public health; and a 'Superbugs Library' and a 'Superbugs CV' providing the academic background of the project.

#### *Teacher brochure*

In March 2023, bilingual printed copies of a professionally designed 2-page brochure were sent to all 1,473 primary and secondary schools in Wales; an electronic version was circulated by email to all secondary schools in Wales and to primary schools in South Wales. As this way of communication only allowed us to reach the school head offices, and it was not clear whether and to what extent the information would be disseminated to relevant staff, we also targeted teachers directly. As such, printed copies were distributed at Cardiff University's "Science in Health Live!" engagement event for year 12 pupils in March 2023 and at the Cardiff University Teacher and Career Adviser's Conference in May 2023. In addition, we identified science, biology and STEM teachers or school departments based on their Twitter biographies, and between March and September 2023 sent them a link to the teacher brochure via direct messaging. From a total of 684 messages sent out, at least 534 were actually opened by the users (marked as 'seen' on Twitter). Encouragingly, from those educators/departments who did see our message, 39 (7.3%) reacted with a 'thumbs up', heart or similar emoji, and a further 54 (10.1%) sent a reply – with messages ranging from a simple "*Thanks!*" or "*Looks nice!*" to more detailed feedback (**Supplemental Table S2**), at times triggering longer conversations and an expansion of our collaborative network (not shown). All in all, the corresponding page of *Superbugs.online* was viewed 740 times, and the PDF version of the brochure was downloaded 123 times, of which seven times (5.7%) in Gaelic and five times (4.1%) in Welsh; note that the Irish translation only became available in September 2023 and thus is not included in these download numbers.

#### *Online traffic analysis*

From its launch until 30 September 2023, *Superbugs.online* attracted a total of 19,155 visits by 16,435 unique visitors, resulting in 33,640 page views. While 62.5% of all visitors were based in the UK as our main target audience, Superbugs had a significant global reach spanning Europe, Asia, Africa, Australia and the Americas (**Supplemental Figure S5**). In parallel to the traffic on the website, we gained 2,018 followers and accumulated >775,000 impressions on Twitter. Most traffic to *Superbugs.online* came from direct sources (64.4%), social media (16.4%) and online searches (113%). A smaller proportion of traffic (7.8%) came via referrals from other websites, including links from Cardiff University; Hwb (Digital Learning for Wales, the Welsh Government's provision of educational tools); Gaelic Education (maintained by Stòrlann); and the Federation of European Microbiological Societies (FEMS). Given the non-descriptive nature of *Superbugs.online* and depending on how visitors find it, not all visits start on the 'Welcome' page. However, the time course of the traffic to the start of the 'Adventure Trail' closely mirrored that on the Welcome page (**Supplemental Figure S3A**), suggesting that approximately every third visit on the 'Welcome' page resulted in a subsequent visit of the 'Adventure Trail'. This agrees with a measured bounce rate of 75.9% of the Welcome page (the percentage of visits that only result in a single view) and an exit rate of 58.5% (the percentage of page views that results in a visitor leaving the site). The Adventure Trail had a bounce rate of 63.0% and with 29.7% a very low exit rate.

Traffic data showed that the 'Adventure Trail' was the most popular part of *Superbugs.online* overall, with 2,493 views of the start page and a further 6,305 views of the individual steps of the adventure trail. This was followed by 'Being a Scientist' (1,585 views), 'Teacher's Corner' (1,323 views), 'Meet the Team' (1,173 views) and the 'Wordle' (1,165 views) (**Supplemental Table S1**). The launch of a multilingual science and health-focused Wordle puzzle – to our knowledge the only one of its kind – in February 2022 led to substantial traffic but did not appear to encourage visitors to explore other content on the website, showing that this game served mainly as a popular stand-alone item and attracted a different audience that was mainly interested in the game itself rather than the scientific content (**Supplemental Figure S3B**). In contrast, the distribution of a bespoke brochure to science teachers and schools in March/April 2023 coincided with considerable views of sections like the 'Adventure Trail' and 'Story Time' (**Supplemental Figure S3B**). As the launch of the Wordle game was heavily promoted on social media it is perhaps not surprising that the web traffic to the corresponding page correlated closely with the total Twitter impressions at the time, demonstrating the effectiveness of Twitter in directing traffic to specific web content (**Supplemental Figure S3C**). Detailed analysis confirmed that at the time of the Wordle launch the majority of the traffic to *Superbugs.online* came indeed via Twitter while the targeted promotion of the teacher brochure by printed leaflets, emails and direct messages led to a spike in direct web traffic (**Supplemental Figure S4**).

**Supplemental Table S1. Major subsections of the Superbugs website.** Unless stated otherwise, all pages are available in English, Welsh, Irish and Scottish Gaelic; languages can be easily changed using a toggle at the bottom of each page. URLs for the English version start with 'www'; URLs for the other languages have the '*www'* replaced with '*cy'* for Welsh (*Cymraeg*), '*ga'* for *Irish* (*Gaeilge*), or '*gd'* for Scottish Gaelic (*Gàidhlig*). Total page views are given for the period from 1 October 2021 until 30 September 2023.

| Description | Live since | URL | Page views |
|---|---|---|---|
| Welcome (English) | 10/2021 | *https://www.superbugs.online* | 7,804 |
| Welcome (Welsh) | 10/2021 | *https://cy.superbugs.online* | |
| Welcome (Scottish Gaelic) | 07/2022 | *https://gd.superbugs.online* | |
| Welcome (Irish) | 08/2023 | *https://ga.superbugs.online* | |
| Adventure Trail, start page | 10/2021 | *…/activities* | 2,493 |
| Behind the Scenes * | 10/2021 | *…/superblog* | 375 |
| Being a Scientist | 10/2021 | *…/being-a-scientist* | 1,586 |
| Bookshelf ** | 03/2023 | *…/books* | 310 |
| Closing Credits | 10/2021 | *…/closing-credits* | 59 |
| Contact Us | 10/2021 | *…/get-in-touch* | 634 |
| Data Protection Privacy Notice | 10/2021 | *…/privacy-notice* | 21 |
| Downloads | 10/2021 | *…/downloads* | 876 |
| Games | 07/2022 | *…/games* | 363 |
| Hall of Fame | 10/2021 | *…/uploads* | 231 |
| Meet the Scientists | 10/2021 | *…/meet-the-scientists* | 140 |
| Meet the Team | 10/2021 | *…/meet-the-team* | 1,173 |
| Participating Schools | 10/2021 | *…/participating-schools* | 312 |
| Pop-up Shop | 10/2021 | *…/pop-up-shop* | 513 |
| Science Fun in Wales and Beyond *** | 10/2022 | *…/science-in-wales* | 119 |
| Spelling Bee ** | 06/2022 | *…/spelling-bee* | 96 |
| Story Time | 10/2021 | *…/reading-corner* | 688 |
| Superbugs International ** | 04/2023 | *…/international* | 168 |
| Superbugs Library | 10/2021 | *…/superbugs-library* | 101 |
| Superbugs Project CV ** | 09/2022 | *…/superbugs-cv* | 54 |
| Teacher Brochure | 03/2023 | *…/superblog/brochure* | 754 |
| Teacher's Corner | 10/2022 | *…/teachers-corner* | 1,323 |
| Wordle | 02/2022 | *…/wordle* | 1,165 |

* Mainly in English and Welsh; some blog entries also in Gaelic and/or Irish

** In English only

*** In English and Welsh only

**Supplemental Table S2. Anecdotal feedback from teachers receiving the Superbugs brochure.** Examples were selected from a total of 54 replies to a personal message with the link to the teacher brochure sent via Twitter direct messaging.

| **Representative replies from teachers, science departments or schools** |
|---|
| *"Amazing! I'll take a look."* |
| *"Sounds interesting, I will have to check them out."* |
| *"I look forward to going through your resources."* |
| *"Thanks a million, will definitely check this out."* |
| *"These seem really fun and interactive!"* |
| *"Looks great, I'll share with the team."* |
| *"I am excited to look through all of the resources and see how I can use them with my students."* |
| *"That is a great resource to have the language option, I'll definitely pass this along to colleagues."* |
| *"Potentially yes. We teach a topic called microbes in our school."* |
| *"Brilliant, will definitely keep this in mind for next year, thank you!"* |
| *"The adventure trail resources look excellent, I will definitely be using them for my year 7 groups."* |
| *"I'd love to avail of the resources, looks great. […} Would love to touch base again when I'm organising these classes."* |
| *"Great I will dig deeper and show you what we end up doing."* |
| *"We are moving onto viruses and bacteria so your material will be useful."* |
| *"I am about to rewrite our immunology course so will definitely have a look!"* |
| *"We are designing our microbiology course this year and I would be so happy to incorporate materials."* |
| *"This looks like a fantastic initiative. I used to work at Cardiff Med School, great to see this outreach."* |
| *"Having them in Welsh is brilliant!"* |
| *"Great to hear you are translating the resources to Irish."* |

**Supplemental Table S3. Anecdotal feedback from pupils on a preliminary version of the Superbugs website.** Examples were selected from 247 pupils completing the feedback form in July 2021, three months before the official launch of *Superbugs.online*. Pupils were from Deri View Primary School (Abergavenny), Tredegarville Church in Wales Primary School (Cardiff), Ysgol Clywedog (Wrexham) and St Alban's Roman Catholic High School (Pontypool). Any typographical errors in the original answers have been kept.

| What is the most interesting thing you have learned from the website? |
|---|
| *“that microbes can kill people and microbes can also be good in food”* |
| *“microbes grow really fast”* |
| *“that when bacteria mutates it gets weaker or stronger”* |
| *“I learnt that if a virus is the size of a tennis ball- then a bacteria would be the size of a school bus!”* |
| *“not all bacteria is harmful”* |
| *“that antibiotics can kill some good and bad bacteria”* |
| *“Imagine one of your cells was the size of a car. Bacteria in comparison would be the size of a book from your school library. A virus, would be the size of a fly.”* |
| *“the most interesting thing I have learnt was that smallpox was the worlds biggest killer in the world”* |
| *“The difference between different micro-organisms especially the jobs and sizes of each.”* |
| *“The most interesting thing I have learnt is that nitrogen is an essential element for our bodies”* |
| *“The thing that I found most interesting was learning about the history of epidemics in the Timeline 'R' Us section and the history of smallpox in the story time section”* |
| *“The black death killed between 75-200 million people in only 4 years”* |
| *“the most intresting thing that i learnt is that bacteria plays an important role in how our planet works.”* |
| *“something interesting I learnt was probably how many different cells there are! There are so many different kinds with different jobs and different looks!”* |
| *“viruses are the smallest cells and you can fit 100 million of them on your thumbnail.”* |
| *“That no matter how hard you try to resist bacteria it will always find a way to come back.”* |
| *“that microbes spread faster than i thought.”* |
| *“Jelly fish are over 500 million years old!”* |
| *“that their have been so many virus outbreaks”* |
| *“small pox was alot like how covid is and how it spreads”* |

**Supplemental Table S4. Anecdotal feedback from pupils on the final version of the Superbugs website.** Examples were selected from 22 pupils of the same year 6 class at an English-medium primary school in Cardiff completing the online feedback form in October 2021, shortly after the launch of *Superbugs.online*. Any typographical errors in the original answers have been retained.

| **What is the most interesting thing you have learned from the website?** |
|---|
| *“smallpox dosent exist any more, so thats one less diesease to worry about”* |
| *“What different bacteria's do to your body”* |
| *“How bacteria works and how there are good and bad bacteria”* |

| **What is your favourite part of the website and why?** |
|---|
| *“My favourite part of the website is that it has great stories with amazing facts”* |
| *“My favourite part of the website is the story time because it's fascinating and gets my brain working.”* |
| *“My favourite part of the website is the part where you get to see what the different bacteria do to the different parts of the body.”* |
| *“how to make your on DNA because i found it really cool”* |

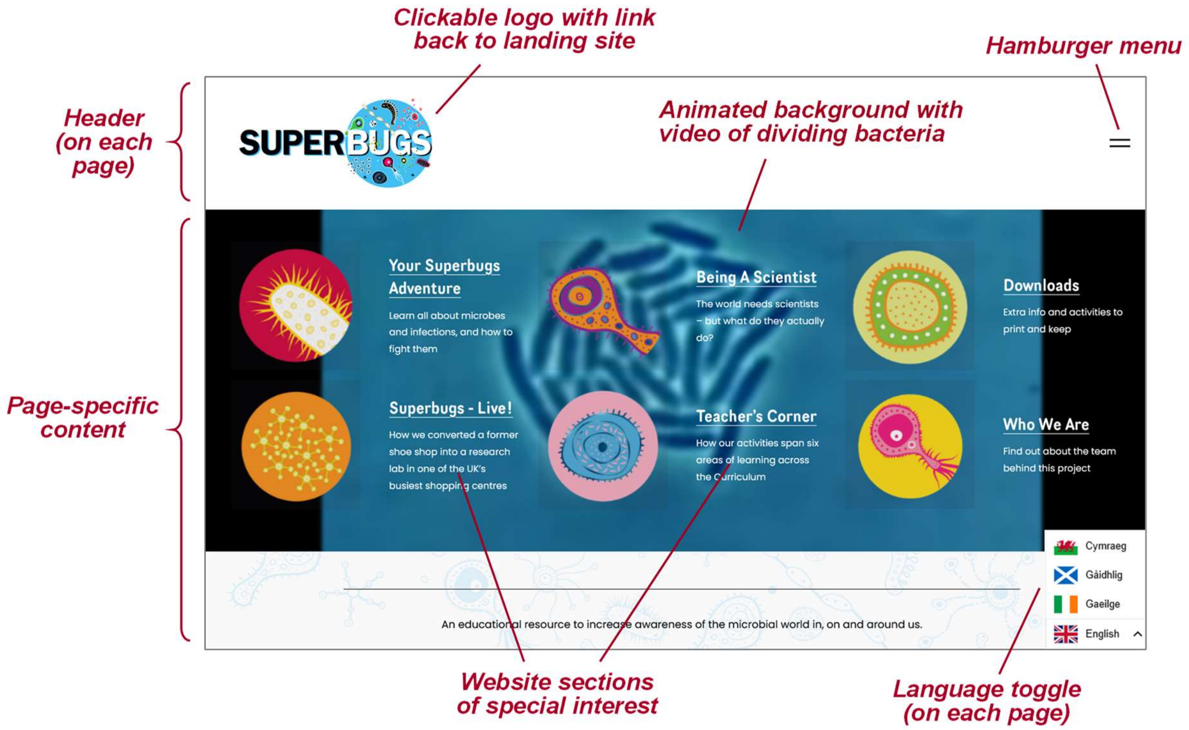


**Supplemental Figure S1**. **Landing page of *Superbugs.online*.** Screenshot taken in September 2023.

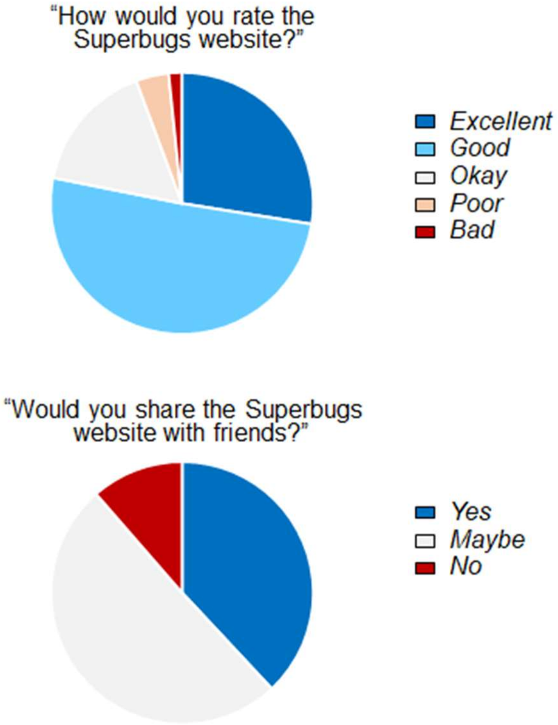


**Supplemental Figure S2. Feedback from 247 pupils attending primary and secondary schools in Wales regarding an early prototype of the Superbugs website.** Pupils were from schools including Deri View Primary School (Abergavenny), Tredegarville Church in Wales Primary School (Cardiff), Ysgol Clywedog (Wrexham) and St Alban's Roman Catholic High School (Pontypool), and completed feedback forms in July 2021, three months before the official launch of *Superbugs.online*.

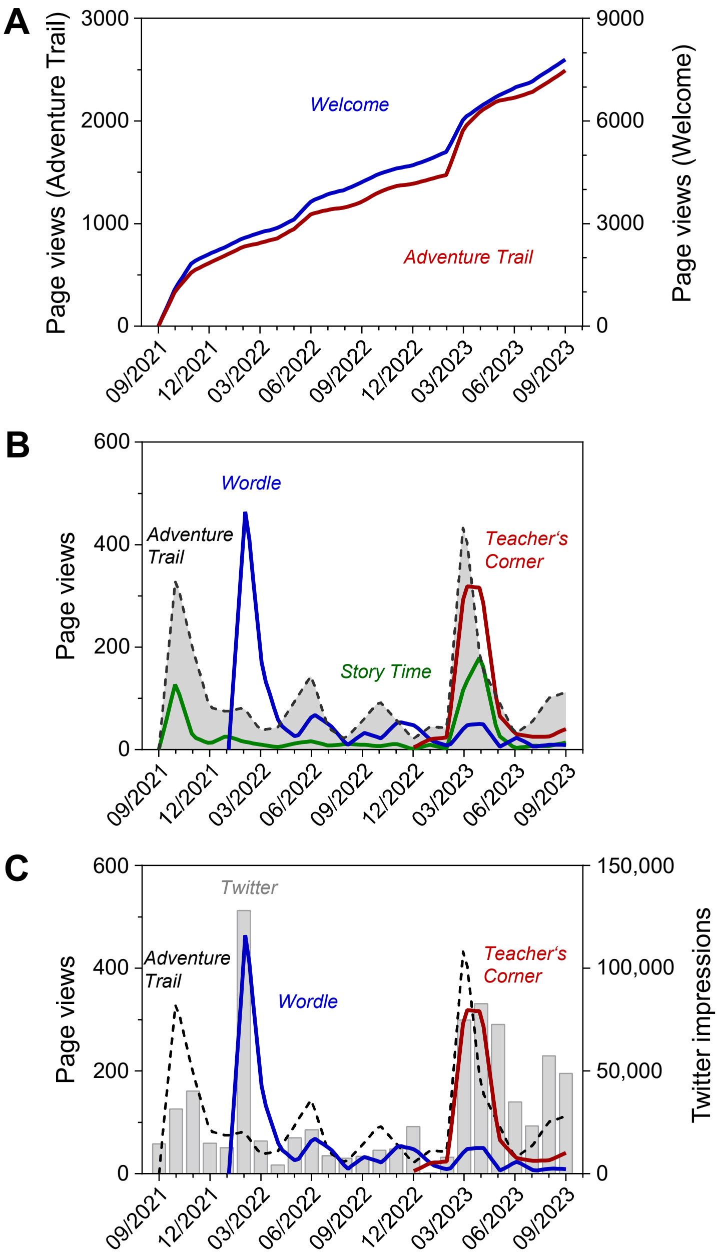


**Supplemental Figure S3. Web traffic to the Superbugs website between 1 October 2021 and 30 September 2023.** ***(A)*** Cumulative traffic to the 'Welcome' page (right-hand scale) and to the 'Adventure Trail', 'Being a Scientist' and 'Story Time' pages (left-hand scale). ***(B)*** Monthly traffic to the 'Adventure Trail', 'Teacher's Corner', 'Story Time' and 'Wordle' pages. ***(C)*** Monthly traffic to the 'Adventure Trail', 'Teacher's Corner' and 'Wordle' pages, overlaid with monthly Twitter impressions. Note that the total web traffic is likely to be underestimated as the analytical data only cover visitors who accept the tracking cookie, and multiple users using the same portal (*e.g.* students in a classroom) may be counted as single visitor. Geography may sometimes reflect the location of the internet provider, not the actual location of the visitor.

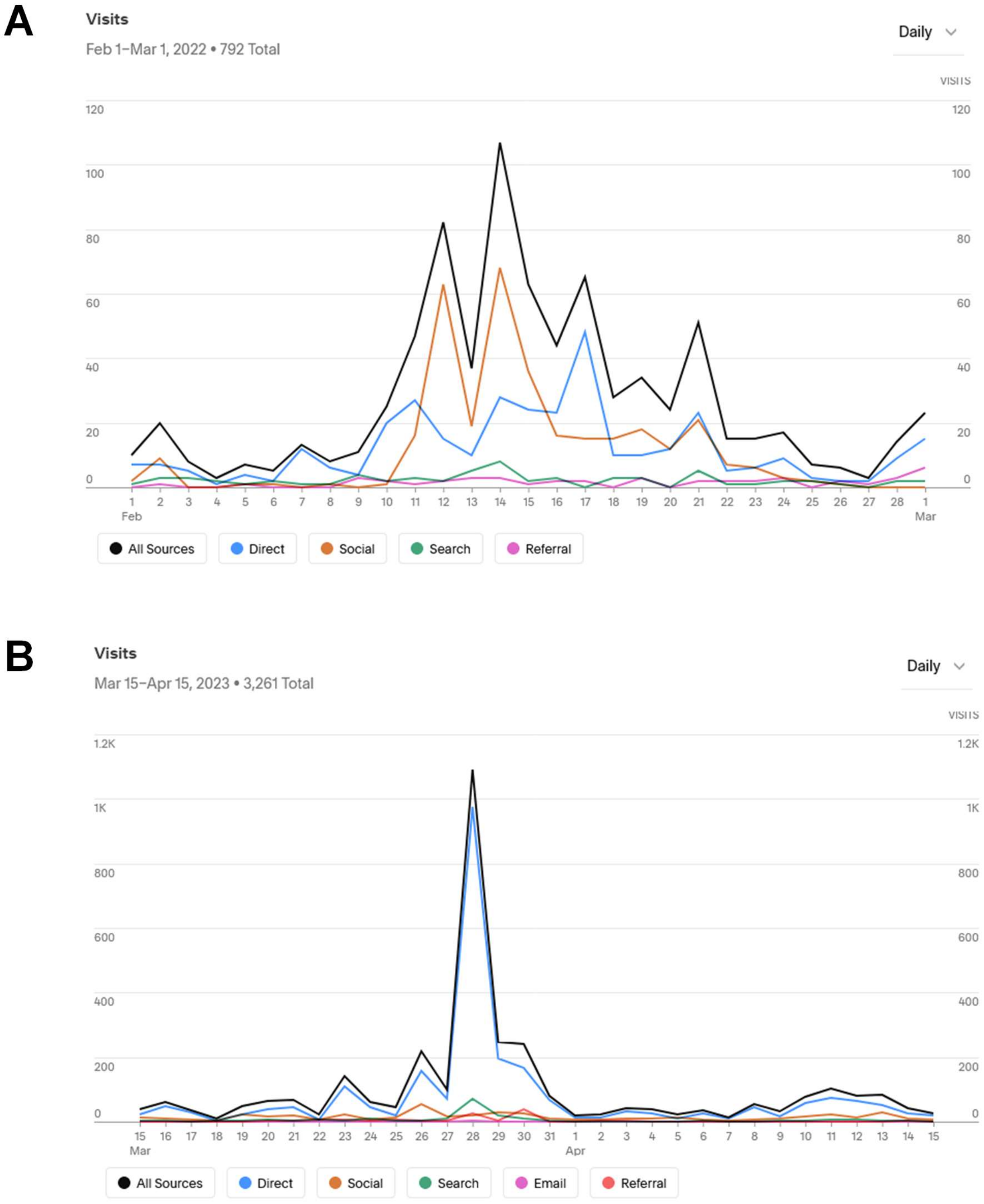


**Supplemental Figure S4. Web traffic to the Superbugs website at the time of the launch of the Wordle in February 2021 *(A)* and the teacher brochure in March/April 2023 *(B)*.** Analysis done using Google Analytics / Squarespace.

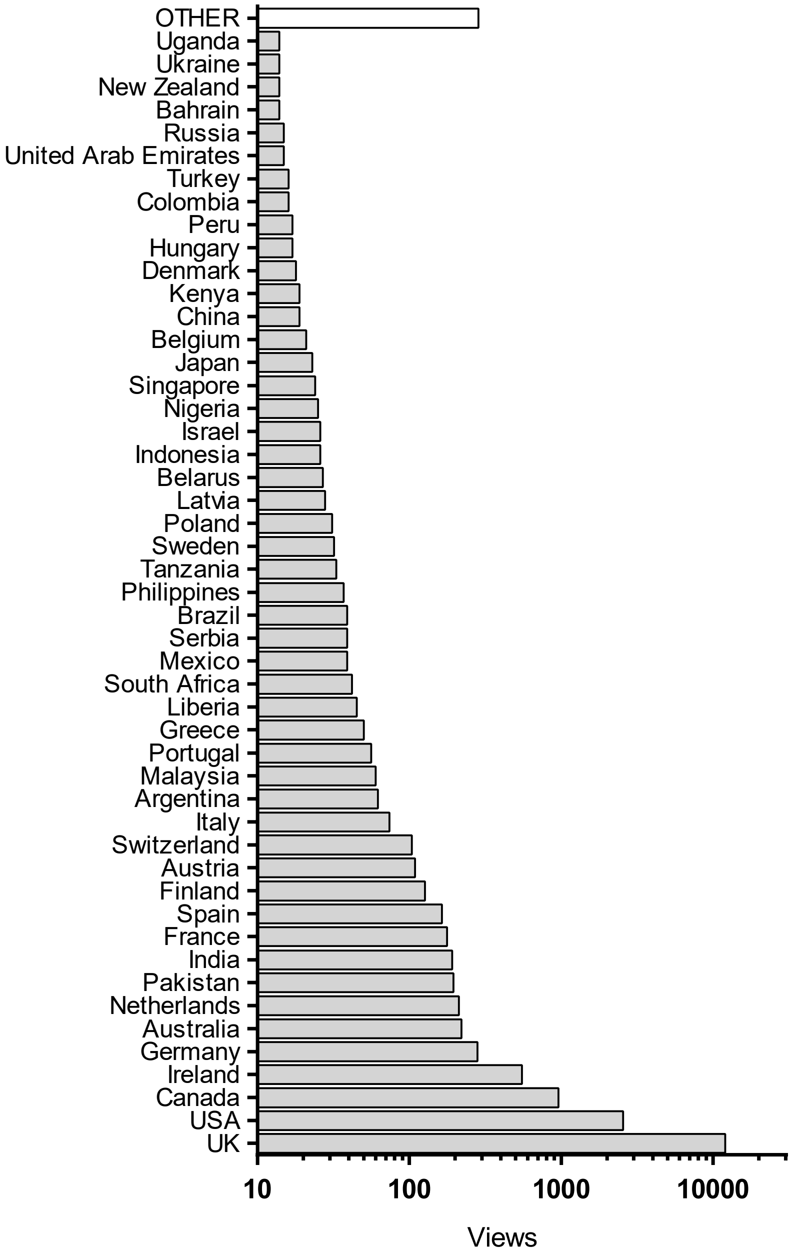


**Supplemental Figure S5. Geographical origin of the visitors of the Superbugs website between 1 October 2021 and 30 September 2023.** Analysis done using Google Analytics / Squarespace.
